# Deep learning-based adaptive detection of fetal nucleated red blood cells

**DOI:** 10.1101/2020.03.06.980227

**Authors:** Chao Sun, Ruijie Wang, Lanbo Zhao, Lu Han, Sijia Ma, Dongxin Liang, Lei Wang, Xiaoqian Tuo, Dexing Zhong, Qiling Li

## Abstract

**Aim:** this study, we established an artificial intelligence system for rapid identification of fetal nucleated red blood cells (fNRBCs).

**Method:** Density gradient centrifugation and magnetic-activated cell sorting were used for the separation of fNRBCs from umbilical cord blood. The cell block technique was used for fixation. We proposed a novel preprocessing method based on imaging characteristics of fNRBCs for region of interest (ROI) extraction, which automatically segmented individual cells in peripheral blood cell smears. The discriminant information from ROIs was encoded into a feature vector and pathological diagnosis were provided by the prediction network.

**Results:** Four umbilical cord blood samples were collected and validated based on a large dataset containing 260 samples. Finally, the dataset was classified into 3,720 and 1,040 slides for training and testing, respectively. In the test set, classifier obtained 98.5% accuracy and 96.5% sensitivity.

**Conclusion:** Therefore, this study offers an effective and accurate method for fNRBCs preservation and identification.

## Introduction

The clinical application of fNRBCs during pregnancy could be classified into two main categories^1,2^. One is the prognosis of possible diseases in pregnant women by counting fNRBCs in umbilical cord blood. Chronic tissue hypoxia results in increased levels of erythropoietin, which, in turn, leads to stimulation of erythropoiesis and increased numbers of circulating nucleated red blood cells (NRBCs)^1,3–5^. Increased umbilical cord levels of erythropoietin have been reported in pregnancies complicated by intrauterine growth restriction, maternal hypertension, preeclampsia, maternal smoking, Rh isoimmunization, and maternal diabetes^5–8^. As expected, each of these conditions has been associated with increased NRBCs in the newborn^9^. The other objective is to screen and extract fNRBCs from maternal peripheral blood for non-invasive prenatal diagnosis (NIPD)^10–12^. The choice of fNRBCs as ideal target cells is based on the following parameters^13,14^: (1) presence of intact nuclei containing the complete fetal genome in fNRBCs, which is a prerequisite for prenatal analysis; (2) limited life span of fNRBCs in the maternal circulation, which can be differentiated morphologically from maternal cells; and (3) presence of distinct cell markers, such as epsilon hemoglobin transferrin receptor (CD71)^15^, thrombospondin receptor (CD36), and glycophorin A (GPA) in fNRBCs that enable isolation of these rare cells from large volumes of maternal blood.

As a result, great attention and research efforts have been devoted to the development of NIPD methods based on circulating fNRBCs. However, the detection of fNRBCs is challenging due to their extremely low concentration against a background predominance of maternal cells (<6 cells per mL; with 109 maternal cells)^16,17^. Several fNRBC enrichment methods based on different principles have been reported, such as density gradient centrifugation (DGC) ^13,18^, fluorescence-activated cell sorting (FACS)^19^, and magnetic-activated cell sorting (MACS)^20^, dielectrophoresis, and microfluidics based technologies^21,22^. Nevertheless, long-term preservation of samples and rapid identification of target cells (fNRBCs) still present considerable challenges.

Since the identification of fNRBCs in large number of cell block slices represent a huge manual burden on pathologists, this field could benefit greatly from an urgent digital revolution^23^. In recent years, the development of computer-aided diagnosis and medical image processing has resulted in emergence of the field of computational pathology^24^. Techniques based on the combination of deep learning and multi-medical specialties has rapidly gained popularity and led to substantial progress in fields such as radiology, ophthalmology, and breast cancer^25–27^. DL-based algorithms have demonstrated remarkable progress in image recognition tasks, in which convolution neural network (CNN) models, as the most prevalent type of deep learning structure, has been reported to surpass human performance^28^, and has become a widely used methodology for analysis in medical imaging^29^.

There are two purposes to the present study. The first is to explore methods suitable for the long-term preservation of fNRBCs. The cell block technique for fixation of fNRBC samples is first introduced. The other objective centers on the establishment of a system based artificial intelligence (AI) to apply supervised learning for the analysis of fNRBC image. Training and validating the CNN model on large-scale data sets is crucial for enhancing the efficiency and accuracy of the model. The fNRBC images were segmented before training, to correct for overestimations. Since it is impossible to rely on expensive and time-consuming manual annotations, we propose a novel adaptive automated region of interest (ROI) extraction algorithm that does not require manual pixel-level annotations. We expect this method to be capable of rapid identification of specific (target) cells in backgrounds cluttered with a large number of maternal peripheral blood, and therefore, to confer simplicity and feasibility to NIPD techniques.

## Materials and Methods

### Ethics statement

This study was approval (XJTU1AF-CRS-2015-001) by the Ethics Committee of the First Affiliated Hospital of Xi’an Jiaotong University. Related informed consent was obtained from the subjects before the study, and all the protocols used were in compliance with the ethical principles for research that involves human subjects of the Helsinki Declaration for medical research^30^.

### Cord blood samples

All umbilical cord blood samples were collected from normal term deliveries (⩾36 weeks). Approximately 9 mL of cord blood from gravidas chosen for this study was collected into anti-coagulant K2-EDTA tubes (BD Vacutainer 366643) containing a proprietary preservative.

### fNRBC enrichment

Blood samples were processed within 2 hours of collection and mononuclear cells were isolated by density gradient centrifugation with Histopaque-1077 (Sigma Chemical, St. Louis, MO, USA), magnetically labelled with anti-CD71 monoclonal antibody (Miltenyi Biotec, Germany), and positively selected by MACS (Miltenyi, Biotec, Germany) according to the protocol provided by the manufacturer.

### fNRBC fixation by cell-block technique

The fNRBC samples were centrifuged at 2000 rpm at normal temperature for 10 min, the supernatant was removed, and the cell-rich layer was collected. Then, the temperature was increased to 40 °C, and the cell-rich layer was absorbed. The samples were transferred to the bottom of the diluted solution (Xi’an Meijiajia, China), loaded into the Li-Shi Thin Prep Liquid-based Cytology and Tissue Embedding Machine (Xi’an Meijiajia, China), and removed after centrifugation at 2000 rpm for 10 min. The samples stood at room temperature (or in a refrigerated room) for 10 min, and were taken out when completely solid. The part without the cell-rich layer were cut off and stored in the embedding box.

### HE staining

Cells were stained with HE staining according to routine protocols^31^. Briefly, after deparaffinization and rehydration, 5 μm longitudinal sections were stained with hematoxylin solution for 3–5 min followed by 5 dips in 1% acid ethanol (1% HCl in 70% ethanol) and rinsed in distilled water. Then, the sections were stained with eosin solution for 5 min, followed by dehydration with graded alcohol and cleaning in xylene. The mounted slides were then imaged using an Olympus BX53 fluorescence microscope (Olympus, Tokyo, Japan). NRBCs were identified by morphology and counted under a light microscope at 400× magnification. Cells were considered as NRBCs if they met the following criteria: diameter was intermediate to that of neutrophils and leukocytes; low nucleus-to-cytoplasm ratio; small, dense, round nucleus; and orthochromatic non-granular cytoplasm.

### Image acquisition and processing

After HE staining, the sections of maternal umbilical cord blood were scanned by a Pathological Section Scanner (Leica SCN 400, Germany). fNRBCs were selected as positive targets, and lymphocytes and neutrophils as negative control targets. The average initial slice size used for ROI extraction was 1,651 × 1,209 pixels.

### Cellular-level ROI extraction

A semi-automated ROI extraction algorithm based on global threshold segmentation and watershed algorithm was proposed^32^.

First, a Gaussian low-pass filter was applied for image pre-processing. To reduce the complexity of ROI extraction, adaptive thresholding methods and mathematical morphology operations were used to segment fNRBCs. Since the HE staining of each fNRBC image was uneven and could fade over time, the adaptive threshold T was calculated by the following formula:

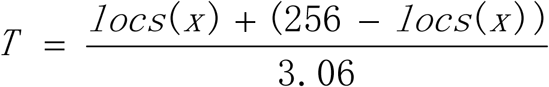

where, the first extreme point on the histogram of the grayscale distribution is denoted as *locs*, and *locs(x)* represents the corresponding abscissa.

Since the global threshold algorithm could not distinguish adjacent cells, we chose the watershed algorithm to detect the single fNRBC. Considering the information in the grayscale image, an improved watershed method based on adaptive thresholding was proposed. First, information on the image gradient was used as prior knowledge, and the watershed algorithm was rendered sensitive to the small extreme line response^33^. Then, the mathematical morphology technique was used to remove cell debris, and over-segmentation was eliminated by bottleneck detection. These steps could reduce the classification burden of the neural network, thereby decreasing the calculation workload. By using effective and robust cellular-level ROI extraction methods, we acquired an accurate cell contour at different magnifications. The experimental results were shown as Figure 1A.

**Figure 1.**
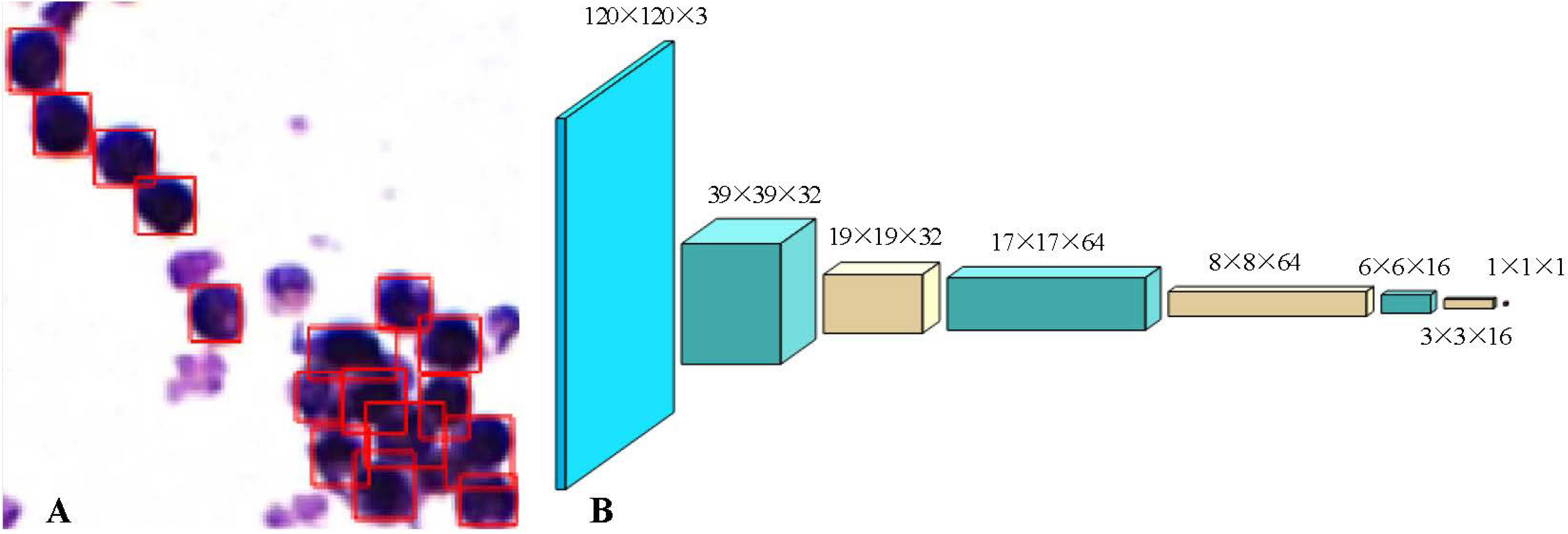
Cellular-level ROI extraction and prediction network. A. An accurate cell contour at different magnifications. B. Schematic representation of the framework of the prediction network.

### Prediction network

We proposed a skillfully-designed network structure p-net to perform classification tasks for fNRBC images. The core of building p-net was to choose the appropriate CNN structure (Supplementary Figure 1) and loss function. VGG-16 had been proven to be successful in the field of medical imaging due to its excellent image feature extraction capabilities. In VGG-16, each input in the layer was linear with the output of the previous layer, resulting in a final output that was a linear representation of the original input. Due to the limitations of linear expression, many features of the original input couldnot be preserved. We needed to combine the data of the input image to generate more features of the image, which would confer greater stability and efficiency to the network. We chose the rectified linear unit (ReLU) function as the activation function.

The p-net was composed of four blocks of convolutional layers, and the final fully connected layers were replaced with a global average pooling layer^34^. Since the size of the ROI patches, an important feature of fNRBCs, was different, we filled the pixels around the ROI patches such that dimensions of 120 × 120 × 3 were achieved, and used these as the input for the network.

The maximum pooling function was chosen as the pooling function to reduce the amount of calculation. Through the training of 10,000 samples, a p-net was fine-tuned on domain-specific dataset. The prediction network framework used in this study was shown in Figure 1B.

### System verification and immunocytochemistry for HbF

HbF was a specific protein found in fNRBCs, which did not exist in maternal erythrocytes and other nucleated cells^35^. Therefore, it could be used as a marker for fNRBC detection. After the establishment of the AI system, some HE sections were selected for system verification, and all positive recognitions were subsequently confirmed by immunocytochemical staining.

Immunocytochemistry studies were performed on 5-μm sections of formalin-fixed, paraffin-embedded tissues. Slides were first deparaffinized and rehydrated. Antigen retrieval was carried out with 0.01 M citrate buffer at pH 6.0. Slides were heated in a 770-W microwave oven for 16 min, cooled to room temper ature, and rinsed in PBS buffer (pH 7.4). The slides were incubated with a 3% hydrogen peroxide solution (hydrogen peroxide: pure water = 1:9) at room temperature in dark for 25 min, followed by washing with PBS buffer (pH 7.4). This step was to block endogenous nonspecific proteins and peroxidase activity. The sections were incubated at 4 °C overnight with HbF (BIOSS, bs-16469r) at a 1:100 ratio (mol). Following a PBS buffer wash, sections were incubated with the HRP-conjugated goat anti-rabbit IgG (G23303, Servicebio, China) at a 1:200 ratio (mol). The sections were then washed and treated with a solution of diaminobenzidine and hydrogen peroxide for 10 min to produce the visible brown pigment. After rinsing, a toning solution (DAB Enhancer, Dako) was used for 2 min to enrich the final color. The sections were counterstained with hematoxylin, dehydrated, and mounted on cover slips with permanent media.

## Results

### Baseline characteristics

This prospective case-control study was conducted at the First Affiliated Hospital of Xi’an Jiaotong University. The study included 4 pregnant women, who delivered a single mature neonate. Table 1 listed the demographic data of these subjects.

**Table 1.** Demographic data of included women

| Patients | Maternal age<br>(years) | Complication | Gestational age<br>(weeks) | Birth weight<br>(g) | Sex of the<br>infant |
| --- | --- | --- | --- | --- | --- |
| 1 | 25 | Gestational<br>diabetes | 37+1 | 3320 | male |
| 2 | 32 | no | 40+4 | 3660 | male |
| 3 | 30 | scar uterus | 39 | 3050 | female |
| 4 | 28 | no | 38+6 | 3760 | male |

### Hematoxylin-eosin (HE) staining of fNRBCs

fNRBCs were detected in the maternal umbilical cord blood. Most fNRBCs were observed to be polychromatic and orthochromatic normoblasts, which was in accordance with the histological features expected of normoblasts (Figure 2A). The single nucleated cell shown by the arrow in Figure 2A represented an fNRBC.

**Figure 2.**
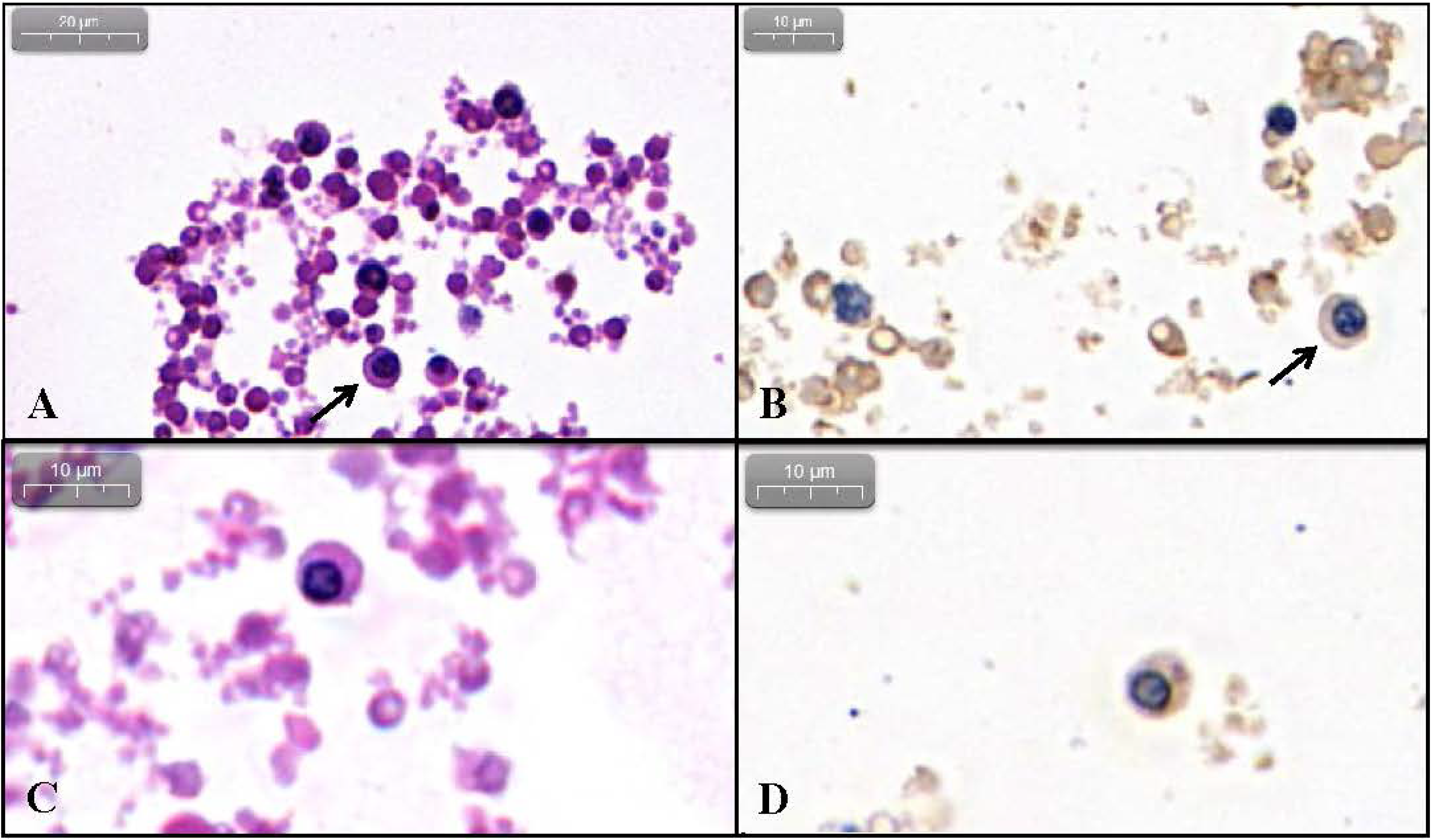
HE staining and HbF immunocytochemical staining of fNRBCs. Panels A and C represented HE staining, while B and D showed cells stained immunocytochemically with HbF. Panels C and D corresponded to the same sample. The single nucleated cells shown by the arrow in A and B represented fNRBCs.

### Hemoglobin F(HbF) immunocytochemical staining of fNRBCs

Blood samples from all 4 patients contained HbF-positive cells. The nuclei were blue stained, while the cytoplasm was not stained (Figure 2B). The single nucleated cell shown by the arrow in Figure 2B represented an fNRBC.

### HE and HbF immunocytochemical staining in serial sections

Eight serial sections (4 μm) were made from each specimen. HE and HbF immunocytochemical staining were performed on odd and even numbered sections, respectively. fNRBCs in the same field of vision were identified based on their distinct morphologies, as observed by HE staining. This was further confirmed by the HbF immunocytochemical staining (Figure 2C, 2D).

### Intelligent identification

We validated the CNN model on a large dataset containing 260 samples, with average slide dimensions of 1,651 × 1,209 pixels (height × width). The total number of positive and negative samples was similar. The training set and test set was randomly split in a 7:3 ratio, based on the original data. Taking the small sample size into account, through data augmentation technology, the original image was flipped to obtain a mirror image, which was rotated by 90°, 180°, and 270°, respectively, thereby expanding the original data set by 8 times. Then, the data set was split into 3,720 and 1,040 slides for training and testing, respectively.

P-net was an end-to-end trainable network, where an image was the input and the result of the threshold operation (a number) represents the output. When the output was close to 1, the sample had a high probability of being positive. Conversely, an output value close to 0 indicates that the sample may belong to the negative group. Since we were more interested in the positive samples with respect to the classification imbalance problem, we chose the precision-recall curve to evaluate the efficiency of the p-net and traditional CNN networks used in this study (Figure 3). Here, Net1 represented a traditional CNN network, while Net2 refered to the network proposed here.

**Figure 3.**
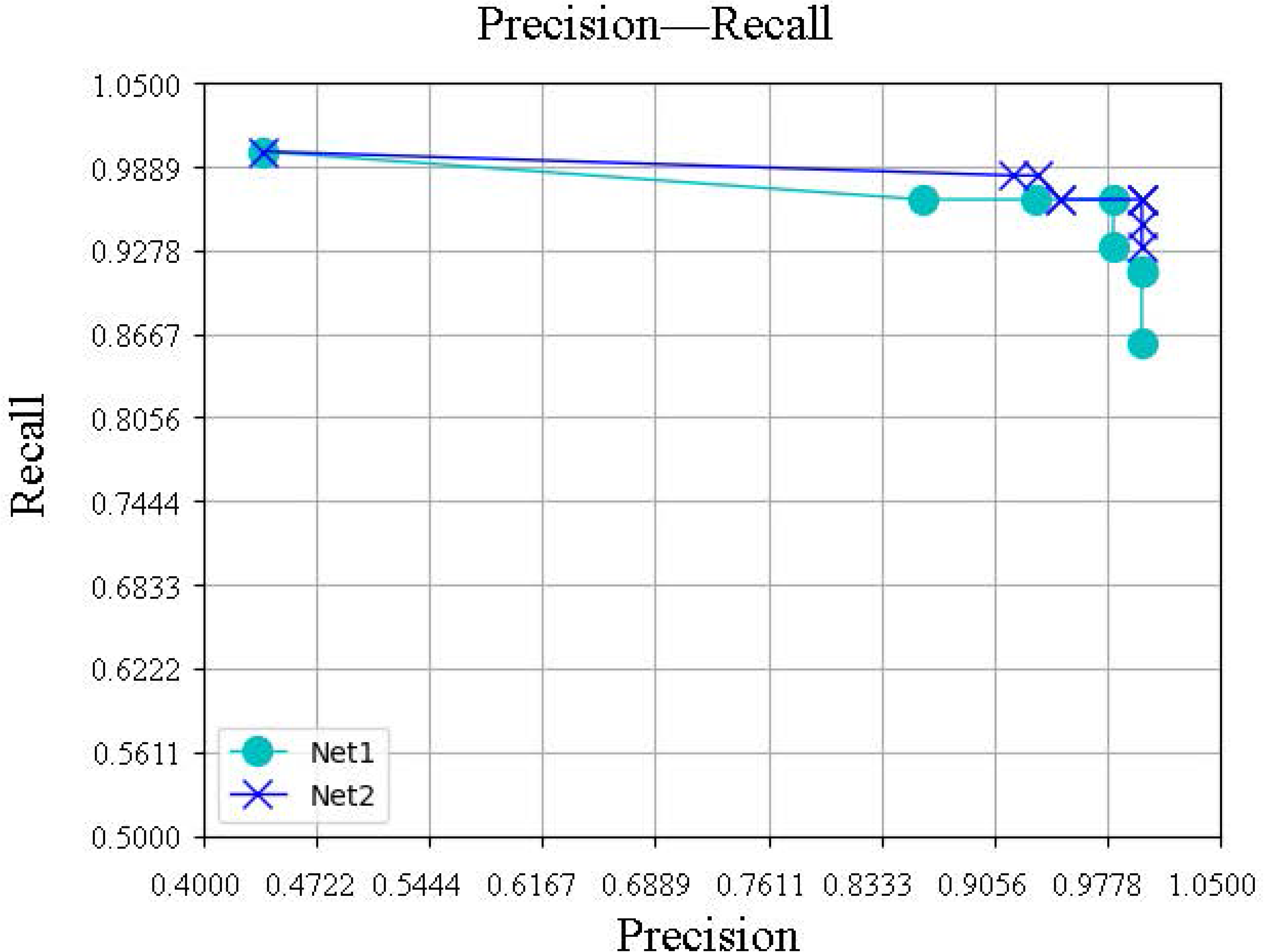
Precision-recall curve of p-net and CNN networks.

The details of the scheme optimization were shown in Table 2. Finally, on the premise of 100% accuracy in the training set, the test set was observed to attain 96.5% sensitivity, 100% specificity, and 98.5% accuracy.

**Table 2.** Comparison of results of different schemes

| Scheme variables | Recognition rate | Sensitivity | Specificity |
| --- | --- | --- | --- |
| Sigmoid activation function | 50% | 100% | 0% |
| The Batch_size parameter is 1 | 79% | 93% | 65% |
| The convolution operators is 5 | 67% | 83% | 51% |
| The output of each layer is pooled | 63% | 65% | 62% |
| Without sample enhancement | 80% | 81% | 79% |
| Our method | 98.5% | 96.5% | 100% |

## Discussion

Prenatal testing based on cell-free DNA (cfDNA) in the maternal plasma has been defined as non-invasive prenatal testing (NIPT) to distinguish it from traditional invasive diagnostic methods such as amniocentesis or chorionic villus sampling^36^. As an excellent screening method deemed acceptable for aneuploidy detection, there are a number of different NIPT platforms, such as massively parallel sequencing, single nucleotide polymorphism, and chromosome-selective sequencing^37^. However, NIPT by itself cannot provide accuracy diagnostics for aneuploidy, and therefore, the karyotyping must be confirmed before or after delivery. Some cfDNA-based techniques called NIPD can provide accurate fetal diagnostic information (thereby offsetting the requirement subsequent confirmation with invasive testing) including fetal sex, rhesus D genotyping, and monogenic disorders^36^. Innovative applications of NIPD, such as digital polymerase chain reaction and next-generation sequencing, have the capability to read more information from cfDNA. Even so, the information cfDNA provides is not as extensive and detailed as that obtained by invasive methods^38^. Meanwhile, there are many challenges in developing NIPD/T services, the most important of which is the content of cfDNA^39^. The latter depends on fetal fraction, and is affected by a variety of factors like gestational age and maternal weight^40^.

Each fetal cell contains all the genetic information of the individual. Therefore, recent studies on NIPD have focused on whole genome sequencing and short tandem repeat identification of fNRBCs and circulating trophoblasts. On the contrary, the rapid selection and separation of target cells still pose considerable challenges to this application^41^. In this study, we report a more effective method for the long-term preservation of fNRBCs. In addition, we have established an AI-based system for the rapid identification of fNRBCs.

Cell block preparations have been used as a complementary technique for increasing diagnostic accuracy in many fields^42^, such as endometrial cytology, malignant pleural effusion, and needle aspiration cytology of thyroid gland^43,44^. In this study, we first proposed a cell block technique for fixation of fNRBC samples. This technique could ensure a uniform distribution of the enriched fNRBCs in the wax block, which is convenient for the identification and isolation of individual fetal cells at a later stage. In addition, the cell slices generated by this technique have no background interference to subsequent immunohistochemistry, fluorescent in situ hybridization, and other molecular pathology assays. fNRBCs are the best target cells for NIPD based on cell block technique. Our method (Cell-Block technique) can not only preserve fNRBCs for a long time, but also facilitate repeated tests using the same sample.

fNRBCs exhibit unique cytological characteristics on HE staining. The nucleus is dense and massive, with the ratio of nucleus to cytoplasm being less than 1/2. There are no granules in the cytoplasm^45^, which is positive stained. Therefore, we used the traditional density gradient centrifugation and MACS (anti-CD71) method to separate fNRBCs from maternal cord blood, and chose conventional HE staining slices for the network-side input. To ensure accurate diagnosis, both HE staining and immunocytochemistry of HbF were used for dual recognition in system verification.

Due to the small number of fNRBCs in maternal peripheral blood, it is not enough for the initial stage of the AI system establishment. In this study, umbilical cord blood of pregnant women was selected as the sample for both input and verification.

Moreover, we introduced artificial intelligence technologies and expect this system to quickly and easily identify fNRBCs in the background predominance of maternal cells. To reduce the complexity in the image classification algorithm, we proposed an adaptive ROI extraction method for fNRBC images. In addition, we have comprehensively utilized the visual information perceived by the network and constructed a novel pathological recognition network, which would have significant contributions in improving the means and methods of non-invasive medical diagnostics.

In general, this report on an AI system of fNRBC identification lays the foundation for subsequent cell collection, sequencing, and prenatal diagnosis. In future, we would conduct further investigations on maternal peripheral blood, and continue to optimize the system, such that it can be devoted to fNRBC detection in NIPD.

---

**Supplementary figure 1.**
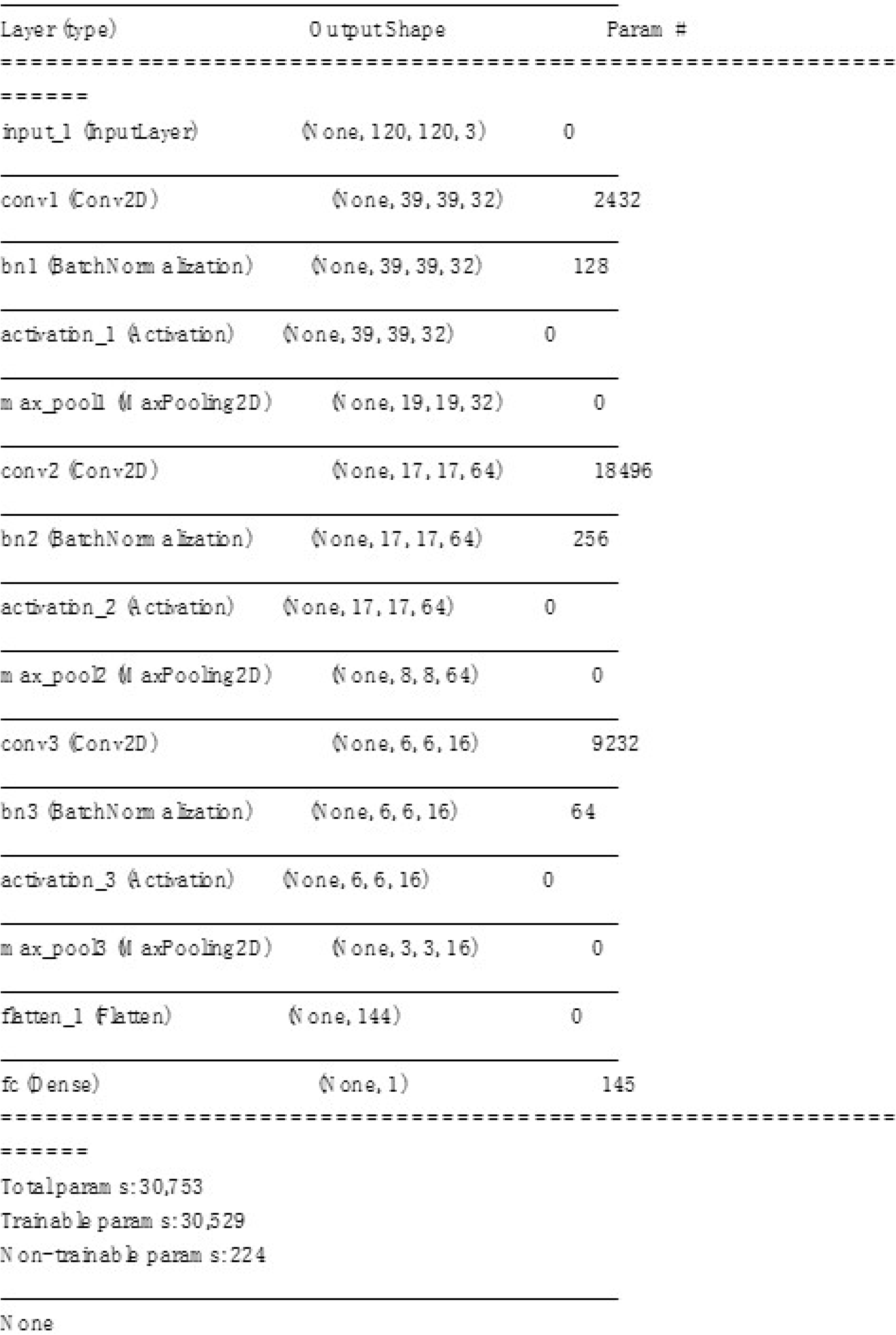
Structure of the CNN model.

## Acknowledgements

This work was supported by the Clinical Research Award of the First Affiliated Hospital of Xi’an Jiaotong University, China (XJTU1AF-2018-017, XJTU1AF-CRF-2019-002), the Major Basic Research Project of Natural Science of Shaanxi Provincial Science and Technology Department (2017ZDJC-11), the Key Research and Development Project of Shaanxi Provincial Science and Technology Department (2017ZDXM-SF-068, 2019QYPY-138), and Shaanxi Provincial Collaborative Technology Innovation Project (2017XT-026, 2018XT-002). The funders had no role in study design, data collection and analysis, decision to publish, or preparation of the manuscript.

## Author contributions

Q.L. and D.Z. conceived and designed the study. C.S., L.Z., L.H. and S.M. performed the laboratory experiments. L.W. finished image acquisition and processing. R.W. analyze and interpret of data. C.S. and R.W. wrote the first draft of the manuscript. D.L. and X.T. revised the manuscript.

## References

1. Kil TH, Han JY, Kim JB, et al. A study on the measurement of the nucleated red blood cell(nRBC) count based on birth weight and its correlation with perinatal prognosis in infantswith very low birth weights. Korean J Pediatr. Feb 2011;54(2):69‐78.

2. Krajewski P,Welfel E, Kalinka J, Pokrzywnicka M, Kwiatkowska M. [Evaluation of therelationship between circulating nucleated red blood cells count and inborn infection inneonates]. Ginekol Pol. Jan 2008;79(1):17–22.

3. Li J,Kobata K, Kamei Y, et al. Nucleated red blood cell counts: an early predictor of braininjury and 2–year outcome in neonates with hypoxic–ischemic encephalopathy in the era ofcooling–based treatment. Brain Dev. Jun 2014;36(6):472–478.

4. Masoudi Z,Akbarzadeh M, Vaziri F, Zare N, Ramzi M. The effects of decreasing maternalanxiety on fetal oxygenation and nucleated red blood cells count in the cord blood. Iran J Pediatr. Jun 2014;24(3):285–292.

5. Boskabadi H,Zakerihamidi M, Sadeghian MH, Avan A, Ghayour–Mobarhan M, Ferns GA. Nucleated red blood cells count as a prognostic biomarker in predicting the complications ofasphyxia in neonates. J Matern Fetal Neonatal Med. Nov 2017;30(21):2551–2556.

6. Constantino BT, Rivera GKQ. Cutoff Value for Correcting White Blood Cell Count for NucleatedRed Blood Cells: What is it? Why is it Important? Lab Med. Oct 10 2019;50(4):e82–e90.

7. Davari–Tanha F, Kaveh M, Nemati S, Javadian P, Salmanian B. Nucleated red blood cells countin pregnancies with idiopathic intra–uterine growth restriction. J Family Reprod Health. Jun 2014;8(2):77–81.

8. Walsh BH, Boylan GB, Dempsey EM, Murray DM. Association of nucleated red blood cells andseverity of encephalopathy in normothermic and hypothermic infants. Acta Paediatr. Feb 2013;102(2):e64–67.

9. Gasparovic VE,Ahmetasevic SG, Colic A. Nucleated red blood cells count as first prognosticmarker for adverse neonatal outcome in severe preeclamptic pregnancies. Coll Antropol. Sep 2012;36(3):853–857.

10. Breman AM,Chow JC, U’Ren L,et al. Evidence for feasibility of fetal trophoblastic cell–basednoninvasive prenatal testing. Prenat Diagn. Nov 2016;36(11):1009–1019.

11. Wei X,Ao Z, Cheng L, et al. Highly sensitive and rapid isolation of fetal nucleated red bloodcells with microbead–based selective sedimentation for non–invasive prenatal diagnostics. Nanotechnology. Oct 26 2018;29(43):434001.

12. Feng C,He Z, Cai B, et al. Non–invasive Prenatal Diagnosis of Chromosomal Aneuploidies andMicrodeletion Syndrome Using Fetal Nucleated Red Blood Cells Isolated by NanostructureMicrochips. Theranostics. 2018;8(5):1301–1311.

13. Mavrou A,Kouvidi E, Antsaklis A, Souka A, Kitsiou Tzeli S, Kolialexi A. Identification ofnucleated red blood cells in maternal circulation: a second step in screening for fetalaneuploidies and pregnancy complications. Prenat Diagn. Feb 2007;27(2):150–153.

14. Redline RW. Elevated circulating fetal nucleated red blood cells and placental pathology interm infants who develop cerebral palsy. Hum Pathol. Sep 2008;39(9):1378–1384.

15. Tao D,Shen Y, Feng X, Chen H. The application of CD71 and Hoechst33258 to staining methodfor sorting fetal nucleated red blood cells in the peripheral blood of pregnant women. Zhonghua Yi Xue Yi Chuan Xue Za Zhi. Oct 2000;17(5):352–354.

16. Xiaoyan X,Hanping C. Fetal nucleated red blood cells in maternal peripheral blood andgestational age. Int J Gynaecol Obstet. Nov 2004;87(2):143–144.

17. Kuo PL. Frequencies of fetal nucleated red blood cells in maternal blood during different stages of gestation. Fetal Diagn Ther. Nov–Dec 1998;13(6):375–379.

18. Kovalak EE,Dede FS, Gelisen O, Dede H, Haberal A. Nonreassuring fetal heart rate patterns and nucleated red blood cells in term neonates. Arch Gynecol Obstet. May 2011;283(5):1005–1009.

19. Sohda S,Arinami T, Hamada H, Nakauchi H, Hamaguchi H, Kubo T. The proportion of fetal nucleated red blood cells in maternal blood: estimation by FACS analysis. Prenat Diagn. Aug 1997;17(8):743–752.

20. Ganshirt D,Smeets FW, Dohr A, et al. Enrichment of fetal nucleated red blood cells from the maternal circulation for prenatal diagnosis: experiences with triple density gradient andMACS based on more than 600 cases. Fetal Diagn Ther. Sep–Oct 1998;13(5):276–286.

21. Zhang H,Yang Y, Li X, et al. Frequency–enhanced transferrin receptor antibody–labelled microfluidic chip (FETAL–Chip) enables efficient enrichment of circulating nucleated red bloodcells for non–invasive prenatal diagnosis. Lab Chip. Sep 11 2018;18(18):2749–2756.

22. Ma GC,Lin WH, Huang CE, et al. A Silicon–based Coral–like Nanostructured Microfluidics to Isolate Rare Cells in Human Circulation: Validation by SK–BR–3 Cancer Cell Line and Its Utility in Circulating Fetal Nucleated Red Blood Cells. Micromachines (Basel). Feb 17 2019;10(2).

23. Hajdu SI. A note from history: microscopic contributions of pioneer pathologists. Ann Clin Lab Sci. Spring 2011;41(2):201–206.

24. Fuchs TJ,Wild PJ, Moch H, Buhmann JM. Computational pathology analysis of tissue microarrays predicts survival of renal clear cell carcinoma patients. Med Image Comput Comput Assist Interv. 2008;11(Pt 2):1–8.

25. Hosny A,Parmar C, Quackenbush J, Schwartz LH, Aerts H. Artificial intelligence in radiology. Nat Rev Cancer. Aug 2018;18(8):500–510.

26. Schmidt-Erfurth U,Sadeghipour A, Gerendas BS, Waldstein SM, Bogunovic H. Artificial intelligence in retina. Prog Retin Eye Res. Nov 2018;67:1–29.

27. Zhou LQ,Wu XL, Huang SY, et al. Lymph Node Metastasis Prediction from Primary BreastCancer US Images Using Deep Learning. Radiology. Nov 19 2019:190372.

28. He K,Zhang X, Ren S, Jian S. Deep Residual Learning for Image Recognition. Paper presented at: 2016 IEEE Conference on Computer Vision and Pattern Recognition (CVPR)2016.

29. Litjens G,Kooi T, Bejnordi BE, et al. A survey on deep learning in medical image analysis. Med Image Anal. Dec 2017;42:60–88.

30. Issue Information–Declaration of Helsinki. J Bone Miner Res. Jun 2018;33(6):BM i–BM ii.

31. Serafini S,Santos MM, Aoun Tannuri AC, et al. Is hematoxylin–eosin staining in rectal mucosal and submucosal biopsies still useful for the diagnosis of Hirschsprung disease? Diagn Pathol. Dec 6 2017;12(1):84.

32. Khan AUM,Torelli A, Wolf I, Gretz N. AutoCellSeg: robust automatic colony forming unit(CFU)/cell analysis using adaptive image segmentation and easy–to–use post–editing techniques. 2018;8(1).

33. Ma H,Beiter R, Gaultier A, Acton ST, Lin Z. OSLO: Automatic Cell Counting and Segmentation for Oligodendrocyte Progenitor Cells. Paper presented at: IEEE International Conference on Image Processing (ICIP)2018.

34. Esteva A,Kuprel B, Novoa RA, et al. Dermatologist–level classification of skin cancer with deep neural networks. Nature. Feb 2 2017;542(7639):115–118.

35. Mavrou A,Kolialexi A, Antsaklis A, Korantzis A, Metaxotou C. Identification of fetal nucleated red blood cells in the maternal circulation during pregnancy using anti–hemoglobin–epsilon antibody. Fetal Diagn Ther. Sep–Oct 2003;18(5):309–313.

36. Skrzypek H,Hui L. Noninvasive prenatal testing for fetal aneuploidy and single gene disorders. Best Pract Res Clin Obstet Gynaecol. Jul 2017;42:26–38.

37. Taylor-Phillips S,Freeman K, Geppert J, et al. Accuracy of non–invasive prenatal testing using cell–free DNA for detection of Down, Edwards and Patau syndromes: a systematic review and meta–analysis. BMJ Open. Jan 18 2016;6(1):e010002.

38. Breveglieri G,D’Aversa E,Finotti A, Borgatti M. Non–invasive Prenatal Testing Using Fetal DNA. Mol Diagn Ther. Apr 2019;23(2):291–299.

39. Jenkins LA,Deans ZC, Lewis C, Allen S. Delivering an accredited non–invasive prenatal diagnosis service for monogenic disorders and recommendations for best practice. Prenat Diagn. Jan 2018;38(1):44–51.

40. Kinnings SL,Geis JA, Almasri E, et al. Factors affecting levels of circulating cell–free fetal DNAin maternal plasma and their implications for noninvasive prenatal testing. Prenat Diagn. Aug 2015;35(8):816–822.

41. Pin-Jung C,Pai-Chi T,Zhu Y, et al. Noninvasive Prenatal Diagnostics: Recent Developments Using Circulating Fetal Nucleated Cells. Curr Obstet Gynecol Rep. Mar 2019;8(1):1–8.

42. Bandyopadhyay A,Bhattacharyya S, Roy S, Majumdar K, Bose K, Boler AK. Cytology Microarray on Cell Block Preparation: A Novel Diagnostic Approach in Fluid Cytology. J Cytol. Apr–Jun 2019;36(2):79–83.

43. Abe H,Takase Y, Sadashima E, et al. Insulinoma–associated protein 1 is a novel diagnostic marker of small cell lung cancer in bronchial brushing and cell block cytology from pleural effusions: Validity and reliability with cutoff value. Cancer Cytopathol. Sep 2019;127(9):598–605.

44. Woo CG,Son SM, Han HS, et al. Diagnostic benefits of the combined use of liquid–based cytology, cell block, and carcinoembryonic antigen immunocytochemistry in malignant pleural effusion. J Thorac Dis. Aug 2018;10(8):4931–4939.

45. Zou L,Ye X, Xu K, Zhu J. Isolation of fetal nucleated red blood cells from maternal blood. J Tongji Med Univ. 2000;20(2):169–171.

